# Transient Naive Treatment (TNT) iPS cells do not feature Sendai virus expression: Response to Sendai virus persistence questions the transient naive reprogramming method for iPSC generation

**DOI:** 10.1101/2024.07.10.602807

**Authors:** Sam Buckberry, Xiaodong Liu, Daniel Poppe, Jia Ping Tan, Geoffrey J. Faulkner, Jose M Polo, Ryan Lister

## Abstract

**Summary:** The Transient Naive Treatment (TNT) reprogramming method involves a brief period of culturing in naive media early in the reprogramming process. Through conducting extensive molecular, epigenetic, and functional analyses on numerous induced pluripotent stem cell lines, we have demonstrated that TNT reprogramming facilitates epigenetic memory erasure and produces induced pluripotent stem (iPS) cells more similar to embryonic stem (ES) cells than conventional reprogramming methods^1^. In 2024, De Los Angeles *et al*. posted a ‘contradictory results’ preprint on *bioRxiv* titled ‘*Sendai virus persistence questions the transient naive reprogramming method for iPSC generation*’^2^. Their claims include: 1) that Sendai virus expression in naive cultured cells questions the TNT reprogramming method, 2) the possible selection of cells (clonal expansion) with Sendai virus expression during TNT reprogramming, 3) Sendai virus genes being expressed in control samples implying a potential sample mix-up. Here, we demonstrate these claims are not directly supported by the data, and emphasize that we assessed the possibility of cell sub-population selection experimentally in Buckberry *et al*. (2023). Moreover, our re-analyses of the data in the context of Sendai virus expression during reprogramming actually highlight the advantages of TNT reprogramming concerning the timing of the naive media treatment and the absence of detectable Sendai virus gene expression.

## Transient-Naive-Treatment (TNT) reprogramming

The TNT method, as originally described, involves a transient 5-day exposure to naive conditions (t2iLGoY media) after an initial 7 days in Fibroblast growth media, with no passaging of cells in naive media (**Fig. 1**). This is in contrast to Naive-iPS cells which are continually passaged in naive media (**Fig. 1**). Thus, the crucial naive media culturing differences between Naive-iPS cells and TNT-iPS cells means that the potential persistence of Sendai virus expression and its associated effects in Naive-iPS cells may not be relevant to TNT reprogramming, as it does not involve extended naive culturing (**Fig. 1**). We emphasize this distinction between Naive-hiPS cells and TNT-hiPS cells, as De Los Angeles *et al*.’s main claims surrounding selection are largely based on data from Naive-hiPS cells.

**Figure 1.**
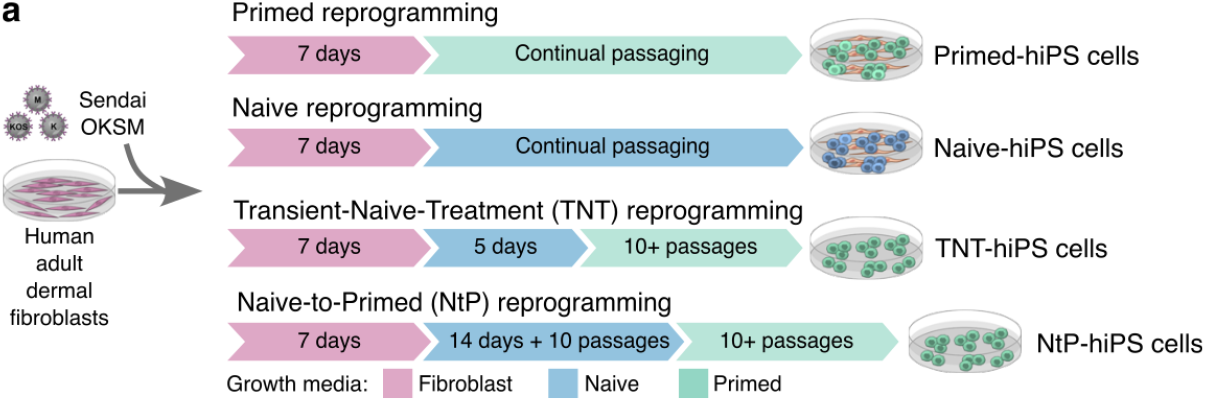
a) Schematic representation of Transient-Naive-Treatment (TNT) reprogramming and Naive-to-Primed (NtP) reprogramming in comparison to conventional Primed and Naive reprogramming, as described in Buckberry *et al*. (2023). Naive media refers to t2iLGoY.

## The data do not indicate that TNT-iPS cells result from preferential selection

In their preprint, De Los Angeles *et al*. state, “*the study utilizes t2iLGoY naive media, which preferentially selects for cells expressing high levels of exogenous reprogramming factors from Sendai virus RNA [*…*]. This selective expression of exogenous factors may significantly contribute to the outcomes observed in the TNT reprogramming*.”

We agree that evaluating any potential selection of cell sub-populations in TNT reprogramming is important. Indeed, we specifically addressed with experiments the potential for clonal selection during TNT reprogramming in Buckberry *et al*. (2023). By using lentiviral insertion mapping with long-read sequencing, we demonstrated that TNT reprogramming does not cause any increase in the rate of clonal selection compared to the conventional Primed reprogramming method (see Buckberry *et al*. (2023) Extended Data Fig. 3i and Supplementary Table 4). In our opinion, it would have been useful and clearer if De Los Angeles *et al*. had acknowledged these experiments, and as a consequence the inconsistency of their postulate with the experimental data from our study.

In formulating their selection hypothesis, De Los Angeles *et al*. draw on their analysis of the Naive-hiPS cells that were continuously cultured in naive media (**Fig. 1**), and show the detection of Sendai virus expression, albeit at levels dramatically lower than those observed in early reprogramming when the highest levels of the Sendai virus is expected (**Fig. 2**). The potential continued expression of the Sendai virus in Naive-hiPS cells has already been reported by us and others in the field^3–5^. However, De Los Angeles *et al*. assert that naive media treatment selects for cells expressing exogenous Yamanaka factors from the Sendai virus and that TNT reprogramming outcomes are affected by cell selection and Sendai virus persistence. We see these assertions as being inconsistent with the experimental design (as explained above) as well as with the analysis they present, which shows that TNT-hiPS cells show no Sendai virus expression or evidence of exogenous Yamanaka factor expression, which should be unexpected if there was selection for cells expressing the Sendai virus. Therefore, our view is that the claims presented by De Los Angeles et al. regarding a selection for cells with exogenous Yamanaka factor expression in TNT reprogramming are not supported by their analysis. Furthermore, our experiments in Buckberry *et al*. (2023) indicate that there is no evidence of increased selection in TNT reprogramming compared to conventional primed reprogramming (see Buckberry *et al*. (2023) Extended Data Fig. 3i and Supplementary Table 4).

**Figure 2:**
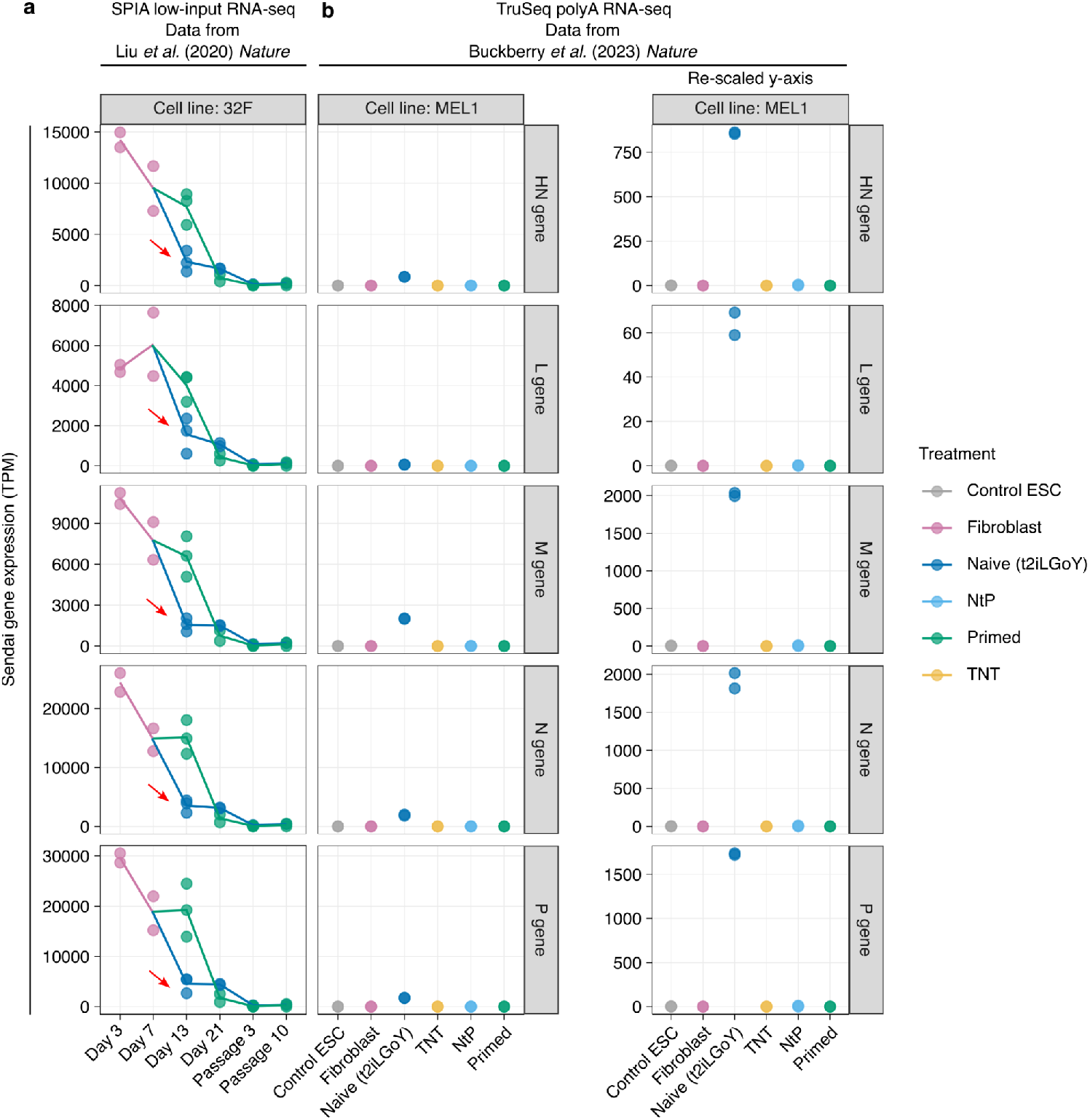
Re-evaluation of the Sendai virus expression estimates generated by De Los Angeles *et al*. **a)** Expression of Sendai virus genes during the time course of reprogramming in Primed and Naive conditions (Cell line: 32F). Red arrows indicate the level of Sendai expression at day 13 of reprogramming is lower in naive reprogramming compared to primed reprogramming. The day 13 timepoint in TNT reprogramming is when cells are transitioned from naive to primed media. **b)** Comparison of hiPS cell reprogramming methods using an isogenic system. Transcripts per million (TPM) expression estimates provided by De Los Angeles *et al*. from their re-analysis of RNA-seq data from Liu et al. (2020) (Cell line: 32F) and Buckberry *et al*. (2023) (Cell line: MEL1). We have only included data in Buckberry *et al*. (2023) to improve data interpretation. Each data point represents an experimental replicate. For the MEL1 cell line, all treatment groups have n=2 replicates.

## TNT-iPS cells do not show Sendai virus expression

To further assess De Los Angeles *et al*.’s claims, we first re-analysed their gene expression estimates (provided by De Los Angeles *et al*.; see Methods) and re-plotted the expression values for each Sendai virus gene across the samples presented in Buckberry *et al*. (2023). De Los Angeles *et al*. presented a conglomeration of RNA-seq data from five different studies using multiple methods and showed an ‘average’ Sendai gene expression for all genes. This included data from cell lines and naive media treatments not used in developing the TNT method or any experiments reported in Buckberry *et al*. (2023). To more clearly visualize the samples and data we used in developing the TNT method, we have replotted De Los Angeles *et al*.’s Sendai virus expression estimates for each gene for the time course of naive and primed reprogramming and our isogenic hES cell and hiPS cell reprogramming experiments. We separated the data for distinct experiment sets and studies to unambiguously show the levels of Sendai virus gene expression (**Fig. 2**). As noted by De Los Angeles *et al*., we observe a reduction of Sendai virus gene expression following infection in fibroblast media, which continues to reduce when cells are transitioned to either primed or naive media during reprogramming (**Fig. 2a**). Importantly, these data actually indicate that at the day 13 reprogramming timepoint in naive media (i.e. after 5 days in naive media), which is the time point at which in TNT reprogramming the cells are then transitioned to primed media, the expression levels of Sendai genes are lower in naive compared to primed media conditions (**Fig. 2a**). Thus, the data actually show the opposite to what would be expected under De Los Angeles *et al*.’s selection model, which posits that culturing in naive media would select for cells with higher Sendai virus expression.

De Los Angeles *et al*. also state that their “*analyses revealed that Sendai genes were expressed in most control PSC samples*”, and refer to their analysis of data from our MEL1 isogenic reprogramming experiments (See Buckberry *et al*. (2023); Fig. 4). Upon closer inspection, we observed that there are negligible or no detected Sendai virus reads in controls (hES, Primed-hiPS, and fibroblast cells), and none detected in TNT-hiPS cells (**Table 1**). This is in contrast to the Naive-hiPS cell samples that have clearly detectable Sendai virus reads (**Table 1**), as expected and as we and others have shown^3,4^. To further evaluate these data, we have re-plotted the raw Sendai virus gene expression estimates De Los Angeles *et al*. generated from our data (**Fig. 2b**). These results show Sendai virus gene expression detected in Naive-hiPS cells, as expected and as we and others have shown previously^3,4^. In contrast, the detection of Sendai virus expression is comparatively extremely low or completely absent in all other experimental groups (**Table 1, Fig. 2b**). Of particular interest, these data actually demonstrate the advantage of the TNT reprogramming system, where after only transient (5-day) exposure to naive conditions early during the reprogramming process, no Sendai virus is detectable in the TNT-hiPS cells. Moreover, these data do not support the claim of “*Sendai genes* [being] *expressed in most control PSC samples*” as their TPM estimates for data from Buckberry *et al*. (2023) has RNA-seq data for 10 pluripotent stem cell samples (iPS or ES cells), and the data show that 6 out of 10 samples have RNA-seq reads mapping to the Sendai genome, with some showing extremely low expression estimates (**Table 1**). We take any claims of sample contamination or potential mix-ups seriously. Claiming that these samples had ‘*significant expression of Sendai virus genes*’, although an overstatement, warrants some analysis of the detection confidence, especially given the low-level signal for all samples except Naive-hiPS cells (**Fig. 1b**), which we investigated further and report the results in the following sections.

**Table 1:** Sendai gene expression as quantified by De Los Angeles *et al*. (2024) *bioRxiv* for data from Buckberry *et al*. (2023).

| Group | Accession | Sendai gene transcripts per million (TPM) |  |  |  |  |
| --- | --- | --- | --- | --- | --- | --- |
|  |  | N gene | P gene | M gene | HN gene | L gene |
| ESC | SRS7483408 | 3.68 | 3.12 | 7.68 | 1.49 | 0.07 |
| ESC | SRS7483409 | 0.13 | 0.13 | 0.16 | 0.04 | 0.00 |
| Fibroblast | SRS7483415 | 0.00 | 0.01 | 0.00 | 0.00 | 0.00 |
| Fibroblast | SRS7483416 | 0.00 | 0.00 | 0.00 | 0.00 | 0.00 |
| Naive (t2iLGoY) | SRS7483410 | 2015.97 | 1721.58 | 2037.59 | 854.11 | 59.00 |
| Naive (t2iLGoY) | SRS7483419 | 8114.79 | 1738.99 | 1994.46 | 862.47 | 69.09 |
| NtP | SRS7483417 | 0.00 | 0.00 | 0.00 | 0.00 | 0.00 |
| NtP | SRS7483418 | 11.52 | 10.72 | 10.78 | 5.64 | 0.33 |
| Primed | SRS7483411 | 0.54 | 0.43 | 0.66 | 0.14 | 0.01 |
| Primed | SRS7483412 | 0.00 | 0.00 | 0.00 | 0.00 | 0.00 |
| TNT | SRS7483413 | 0.00 | 0.00 | 0.00 | 0.00 | 0.00 |
| TNT | SRS7483414 | 0.00 | 0.00 | 0.00 | 0.00 | 0.00 |

## Sendai virus quantification with targeted TaqMan assays demonstrates that Sendai virus is not detected in the control stem cell lines and samples

As explained above, we acknowledge the data from the multiplexed RNA-seq experiment from our study does indicate the presence of some reads that align to the Sendai virus genome, albeit with extremely low levels of Sendai-virus-mapping RNA-seq reads in an uninfected hES cell sample at levels orders of magnitude lower than those observed in the Naive-hiPS cells (**Table 1**). However, this is not direct evidence of a sample mix-up or Sendai virus infection of hES cells.

To investigate further, we performed a series of Sendai virus detection experiments using the specifically designed TaqMan iPSC Sendai Detection Kit (manufactured and validated by Thermo Fisher Scientific as the recommended assay for detecting Sendai virus residue). We profiled our 1) existing cell lines, 2) replicate frozen cell pellets and 3) purified RNA from the exact samples from Buckberry *et al*. (2023). These results show that Sendai virus and exogenous Yamanaka factor expression is only observed in Naive-hiPS cell samples, with no evidence to support Sendai virus infection in hES cells or TNT-hiPS cells (**Fig. 3**). As the origin cell lines, cell pellets, and extracted RNA were all Sendai virus-negative, we can confidently conclude that these hES cell samples were not infected with the Sendai virus.

**Figure 3.**
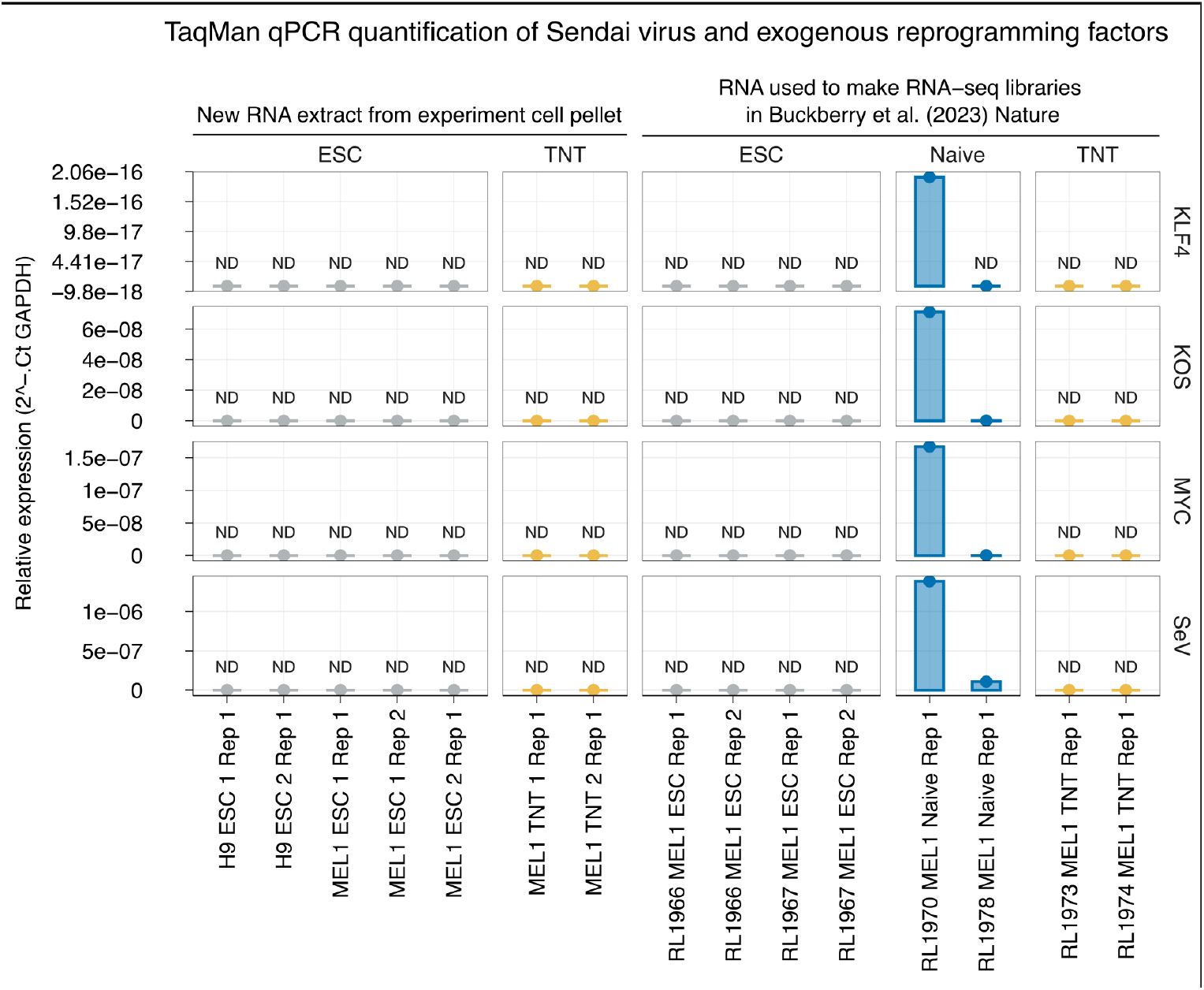
TaqMan qPCR quantification of Sendai virus and exogenous reprogramming factor expression confirms that Sendai virus genes are not detectable in ES cell controls. TaqMan probe-based qPCR quantification of Sendai virus and exogenous Yamanka factor expression in RNA used for the preparation of the RNA-seq libraries (right panels) or replicate frozen cell pellets from hES cells and hiPS cells used in Buckberry *et al*. (2023) (left panels). Y-axis units are relative expression values using the 2^-ΔCt method with GAPDH as the reference gene.

Cross-contamination in multiplexed RNA-seq experiments is widely acknowledged (e.g. ^6^), and can occur from sources such as index-hopping and low-level cross-contamination during library preparation. Our records show that Naive-hiPS and hES cell RNA-seq libraries were prepared at the same time and were processed directly adjacent to each other during preparation, centrifugation, and other processes. Thus, given the very high abundance of Sendai virus RNA molecules in Naive-hiPS cell samples (**Fig. 2; Table 1**), the most plausible explanation for the extremely low hES cell Sendai virus signal is due to some very low-level cross-over contamination or background noise rather than actual viral infection that would drive Sendai virus expression in hES cells.

## Spike-in RNA control analysis indicates that the Sendai virus is not confidently detectable in cell lines other than the Naive-hiPS cells

Given the extremely low abundance of RNA-seq reads mapping to the Sendai virus genome for hES cell samples, and the opportunity for errors and artifacts that could be introduced after RNA extraction, we re-analysed our RNA-seq data with strict mapping and filtering criteria (uniquely mapped read pairs that overlap features by >75%; see **Methods**). To better understand how confident the Sendai virus detection is in these samples, we then performed a limit of detection analysis by leveraging the ERCC spike-in control RNAs we used in the RNA-seq library preparations for these experiments. This analysis shows that the number of reads mapping to Sendai virus genes for non-Naive-hiPS cells is orders of magnitude lower than for Naive-hiPS cells (**Table 2**), and that Sendai virus transcript levels in the control hES cell samples are near the limit of detection based on empirical data from ERCC spike-in transcripts (**Fig. 4**). These data indicate that Sendai virus genes in Naive-hiPS cells are expressed at the higher-end of the dynamic range of expression quantification, whilst Sendai virus genes for hES cells are not all confidently within the bounds of the limit of detection. For example, only 2 of 5 Sendai genes were confidently detectable (>1 TPM) for one hES cell sample, and 0 of 5 Sendai genes for the other hES cell sample (**Fig. 4**). This suggests the hES cell sample was not expressing the full Sendai virus. Therefore, it is highly unlikely these cells were infected by the Sendai virus, and thus the cells would not be expressing exogenous Yamanaka factors.

**Table 2:** Number of RNA-seq fragments (and percentages) assigned to major feature classes.

| Library | Group | Total assigned fragments | Gencode genes | Transposable elements | ERCC controls | Sendai virus genes |
| --- | --- | --- | --- | --- | --- | --- |
| RL1966 | ESC | 17,673,153 | 17,334,595<br>(98.08%) | 250,438<br>(1.42%) | 87,624<br>(0.50%) | 496<br>(0.00281%) |
| RL1967 | ESC | 22,266,164 | 21,867,694<br>(98.21%) | 261,304<br>(1.17%) | 137,145<br>(0.62%) | 21<br>(0.00009%) |
| RL1973 | TNT | 30,140,597 | 29,583,714<br>(98.15%) | 413,000<br>(1.37%) | 143,883<br>(0.48%) | 0<br>(0.00000%) |
| RL1974 | TNT | 25,958,450 | 25,507,596<br>(98.26%) | 327,541<br>(1.26%) | 123,313<br>(0.48%) | 0<br>(0.00000%) |
| RL1977 | NtP | 16,157,584 | 15,840,654<br>(98.04%) | 203,964<br>(1.26%) | 112,966<br>(0.70%) | 0<br>(0.00000%) |
| RL1978 | NtP | 16,092,874 | 15,772,418<br>(98.01%) | 219,108<br>(1.36%) | 100,080<br>(0.62%) | 1,268<br>(0.00788%) |
| RL1970 | Naive | 22,706,181 | 22,011,978<br>(96.94%) | 429,637<br>(1.89%) | 106,802<br>(0.47%) | 157,764<br>(0.69481%) |
| RL1979 | Naive | 15,943,127 | 15,417,178<br>(96.70%) | 341,123<br>(2.14%) | 69,550<br>(0.44%) | 115,276<br>(0.72305%) |
| RL1971 | Primed | 19,443,036 | 18,869,913<br>(97.05%) | 438,262<br>(2.25%) | 134,771<br>(0.69%) | 90<br>(0.00046%) |
| RL1972 | Primed | 25,611,852 | 24,975,886<br>(97.52%) | 512,811<br>(2.00%) | 123,155<br>(0.48%) | 0<br>(0.00000%) |
| RL1975 | Fibroblast | 23,506,597 | 23,282,285<br>(99.05%) | 112,641<br>(0.48%) | 111,671<br>(0.48%) | 0<br>(0.00000%) |
| RL1976 | Fibroblast | 22,525,740 | 22,305,634<br>(99.02%) | 94,058<br>(0.42%) | 126,048<br>(0.56%) | 0<br>(0.00000%) |

**Figure 4.**
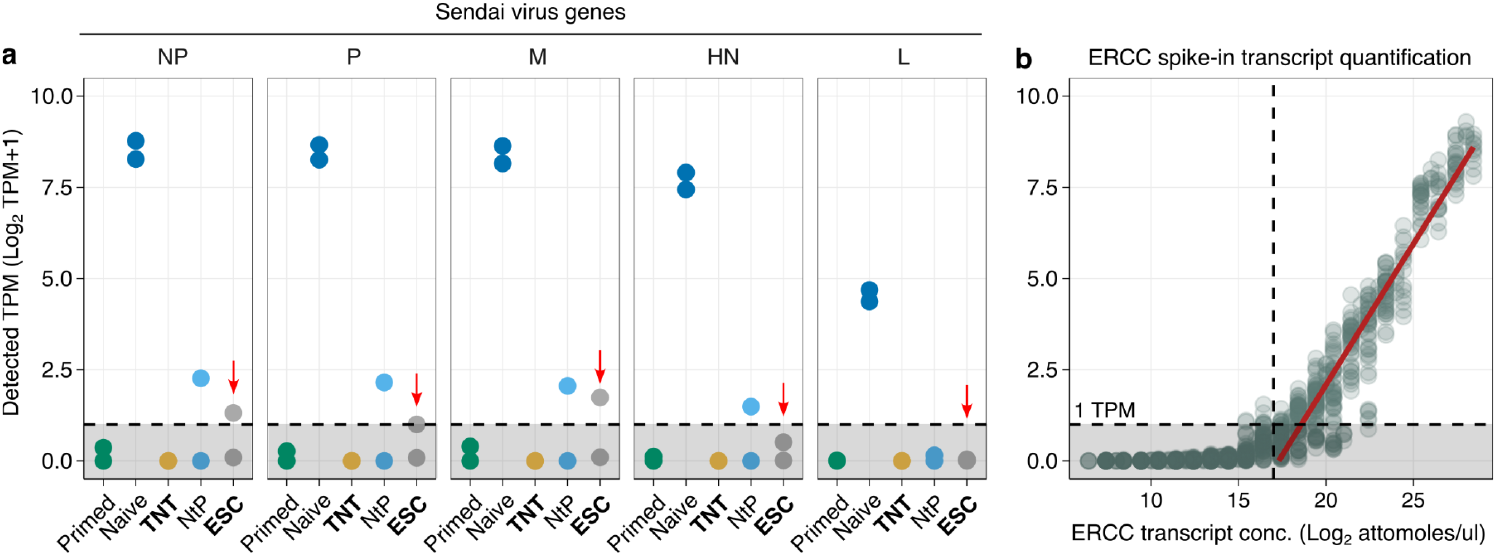
Sendai virus gene quantification coupled with ERCC spike-in RNA dynamic range and limit of detection analysis indicates that only two out of five Sendai genes for only one hES cell sample can be confidently detected. **a)** Sendai gene quantification for MEL1 isogenic pluripotent stem cells in comparison to **b)** Detectable expression range as determined by ERCC spike-in RNA internal controls for all libraries. The plot shows the relationship between the known RNA molecule concentration and the estimated transcript abundance as transcripts per million (TPM) on the log_2_ scale with a linear model fit. Each point on the plot (b) represents an ERCC spike-in transcript. Note that the relationship between transcript concentration and TPM only becomes linear when greater than ∼17 on the x-axis (vertical dashed line), indicating a low confidence of detection around this value. Additionally, the y-axis log_2_ scale emphasizes lower levels of expression around the limit of detection. The horizontal dashed line corresponds to 1 TPM, which these data indicate is a reasonable quantification limit of detection, as below this threshold the confidence in expression quantification is low. Red arrows highlight hES cell Sendai virus detection, indicating that, for hES cells, only one sample has two of five genes that can be confidently detected above 1 TPM. This suggests this hES cell sample was not expressing the full Sendai virus, and therefore, it is highly improbable it was infected.

## TNT-hiPS cell samples show evidence of reprogramming that rules out sample mix-ups with hES cell samples

De Los Angeles *et al*. speculate that the detection of Sendai virus genes in hES cell samples could be the result of a “*mix-up of hESC and TNT hiPSC samples*”. As shown above, we did not detect any Sendai virus or exogenous Yamanaka factor expression in hES cells by the recommended TaqMan assays applied to the original RNA (**Fig. 3**), and the Sendai gene RNA quantification is not confidently detected in the RNA-seq data we present in Buckberry *et al*. (2023) (**Fig. 4**). These data indicate that the most likely explanation is an extremely low level of spurious library contamination at the preparation step.

Nevertheless, we reasoned a way to further rule out this speculated sample mix-up. In Buckberry *et al*., we showed that correction of gene expression with TNT reprogramming, whilst highly effective, is still not complete. In other words, a few genes remain aberrantly expressed in both TNT-hiPS cells and other reprogrammed cells compared to hES cells (See Buckberry *et al*. (2023); Fig. 4c,d). To further demonstrate that De Los Angeles *et al*.’s claim of a potential sample mix-up between the hES cell and TNT-hiPS cell samples is false, here we show examples of the expression levels of such ‘uncorrected’ genes in TNT-hiPS cells, where the expression is not detectable in hES cells, and is detectable in other reprogrammed cells, such as the Primed-hiPS cells (**Fig. 5**). Therefore, if TNT-hiPS cells and hES cells had been mixed up, then one would expect hES cells to show expression signatures of reprogramming. However, this is not what we observe. Together, these data refute the claim that TNT-hiPS cell samples were mixed up with hES cell samples.

**Figure 5.**
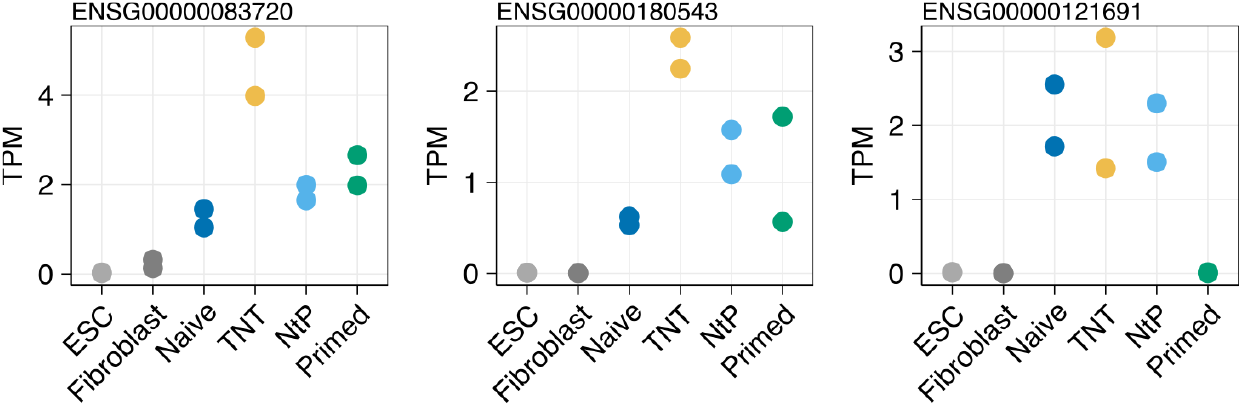
Not all gene expression differences caused by reprogramming are corrected by TNT reprogramming. Plots show RNA-seq expression estimates as transcripts per million (TPM) for an example set of genes not corrected by TNT reprogramming. Points show replicate isogenic hES cell, fibroblast, and derived hiPS cells (n=2 per group). Data are from isogenic MEL1 isogenic reprogramming experiments in Buckberry *et al*. (2023).

## Conclusion

We find De Los Angeles *et al*.*’*s central argument that ‘*Sendai virus persistence questions the TNT method’* is unfounded as the TNT-iPS cell lines do not have any detectable Sendai virus. Their argument against using naive media in reprogramming is not valid for the TNT method due to the ‘transient’ aspect of the method, where cells are not passaged in naive media. Regarding the claim of selection, in Buckberry *et al*. (2023), we showed experimentally that TNT-hiPS cells do not show increased clonal selection compared to conventional primed reprogramming (Primed-hiPSC cells). Finally, De Los Angeles *et al*.’s suggestion of “*a mix-up of hESC and TNT hiPSC samples*” is an overreach, and they provide no evidence as to why those two sample groups would be mixed up. The most plausible explanation for the extremely low levels of Sendai virus RNA-seq reads in hES cells is potential “noise level” signal near the limit of detection, or an unfortunate spurious low-level contamination at the library preparation step or beyond, which is further supported by TaqMan assays showing no Sendai virus RNA in the hES cells or TNT-hiPS cell RNA samples or replicate cell pellets. For these reasons, it is not plausible that hES cells were infected by the Sendai virus, and there would have been no impact on the biology of the cells or other assays in our study.

Importantly, this re-analysis reiterates the benefits of the short transient naive media exposure used in TNT reprogramming, as the expression levels of Sendai genes are actually lower in naive compared to primed media at the day-13 time point when TNT cells are transitioned from naive media to primed media (**Fig. 1a**). The evidence we present here, and the extensive epigenomic and functional analyses presented in Buckberry *et al*. (2023), underscores the advantages of TNT reprogramming, which does not involve extended culturing or passaging in naive media.

## Methods

### De Los Angeles et al. data re-analysis

TPM estimates shown in **Table 1** and **Fig. 2** were generated by De Los Angeles et al. and obtained through the following links: https://github.com/labsyspharm/tnt-rebuttal https://www.synapse.org/Synapse:syn53061839/files/

### polyA RNA-seq (as reported in Buckberry *et al*. 2023)

RNA was extracted using the Agencourt RNAdvance Cell v2 (Beckman Coulter) system following the manufacturer’s instruction with one additional DNAse (NEB) treatment step. RNA amounts and RINe scores were assessed on a TapeStation using RNA Screen Tape (Agilent), and 500 ng of total RNA were used per sample to generate RNA-seq libraries. ERCC ExFold RNA Spike-In mixes (Thermo Scientific) were added as internal controls. Libraries were prepared using the TruSeq Stranded mRNA library prep kit (Illumina), using TruSeq RNA unique dual index adapters (Illumina). Libraries were quantified by qPCR on a CFX96/C1000 cycler (Bio-Rad), multiplexed and sequenced on a NovaSeq 6000 (Illumina) in 2× 53-bp paired-end format.

### Reference genomes and RNA-seq read alignment

Reference genomes GRCh38, NC_075392.1 and ERCC spike-in sequences, along with gencode v27 primary assembly annotations, transposable element annotations, and Sendai gene models were used to create a STAR alignment index.

FASTQ files were processed with `fastp` with the options `--detect_adapter_for_pe --trim_poly_g --trim_front1 1 --trim_tail1 1 --correction`. Reads were aligned with the STAR default options. Post-alignment BAM files were filtered for unique alignments using `samtools -q 255` as STAR uses the MAPQ score of 255 to mark unique alignments. PCR duplicates were marked with `samtools markdup`.

### Expression quantification

Given the goal of confident virus detection, we took a conservative approach to assigning reads to features. These features include gencode genes, transposable elements, Sendai virus genes, and ERCC spike-in transcripts. Quantification was performed with featureCounts with the options `featureCounts -p -T 16 -s 2 --fracOverlap 0.75 --primary --ignoreDup –countReadPairs -B -C -a`. This conservative approach only counts reads that are 1) primary alignments (actually already filtered for with unique alignments), 2) not PCR or optical/exAmp duplicates, 3) both pairs are mapped and not discordant, and 4) reads that overlap the feature being counted by at least 75%.

### Sendai virus quantification by TaqMan PCR

RNA was extracted from cells using RNeasy micro kit (Qiagen) and QIAcube (Qiagen) according to the manufacturer’s instructions. Reverse transcription was then performed using QuantiTect reverse transcription kit (Qiagen). The qPCR assays for detecting KLF4 (Mr04421256_mr), KOS (Mr04421257_mr), MYC (Mr04269876_mr) and SeV (Mr04269880_mr) was subsequently set up using TaqMan™ Gene Expression Assay (FAM) (ThermoFisher Scientific) according to the manufacturer’s instructions. GAPDH (Hs02758991_g1) was used to calculate the relative expression of each assessed gene.

## Data and code availability

https://github.com/SamBuckberry/tnt-biorxiv-response

